# The aging genome exhibits organized vulnerability to somatic mutations

**DOI:** 10.64898/2026.05.21.726885

**Authors:** Joseph Ehlert, Ronald Cutler, Jonah Spector, Bnaya Gross, Orr Levy, Jan Vijg, Xiao Dong, Albert-László Barabási

## Abstract

Somatic mutations accumulate throughout life and have been hypothesized to drive organismal decline. Yet whether these mutations are distributed randomly or whether cells shield their most critical components has remained unresolved. Here we analyze over a million somatic mutations across thirteen human tissues, finding that the aging genome exhibits organized vulnerability, captured by the existence of hypo-mutated genes and longevity-associated pathways that have significantly lower mutation burden. Highly connected network hubs are systematically protected from mutation, while peripheral, condition-specific genes accumulate disproportionate burdens. We show that this organized vulnerability arises from the interplay of two independent mechanisms: transcription-coupled repair, and selective filtering. Finally, we validate our findings under experimental mutagenesis, demonstrating intrinsic mechanisms of protection rather than tissue-specific confounders. These findings reframe the somatic mutation hypothesis: organismal decline may not reflect total mutational burden, but where those mutations fall within the cellular network.

## Introduction

In 1959, Leo Szilard proposed that somatic mutation accumulation drives aging [1], establishing a framework that has since linked genetic changes to multiple age-related pathologies [2, 3, 4]. The emergence of single-cell sequencing has sharpened and expanded Szilard’s hypothesis [5, 6], revealing that mutations accumulate in essentially all organs and tissues in both mice and humans [7, 8, 9]. Moreover, the inverse correlation between mutation rates and lifespan indicates that organisms that accumulate mutations at a lower rate live longer [10, 11]. More direct causal evidence comes from progeroid syndromes, in which germline defects in DNA repair pathways lead to dramatically accelerated aging [12, 13, 14, 15]. While the mutation accumulation hypothesis has driven research on aging for decades, the lack of systematic data has left a fundamental question unresolved: do mutations accumulate randomly across the genome, or does the cell systematically protect its most critical components? The possibility that mutational rates can vary within the same organism is supported by recent measurements: on one end, the observed mutation rates are 1-2 orders of magnitude higher in somatic cells compared to germline, and lower in stem cells than in differentiated cells [16, 17, 18, 19, 20, 21, 22]. Most important, mutation rates vary dramatically across human tissues—from 2.4 mutations/year in brain neurons to 56/year in appendix intestine [23]—reflecting differential mutagen exposures [24, 25], mitotic activity [26], selective constraints [27, 28], and DNA repair efficiency [29, 30].

This heterogeneity raises a deeper question: if mutation rates vary markedly between tissues and organisms, do they also vary systematically across genomic regions within cells? Sporadic evidence suggest that it does: specific mutational signatures have been identified for mutagenic exposures [31], aging [32], and DNA repair activities [33]. Yet, whether mutation variation coincides with regions encoding biological networks within the genome remains unknown.

Here we address the question by relying on the largest available record of aging-related somatic mutations, capturing 1.3 million mutations in 4,684 cells across 13 human tissues [34]. We show that the distribution of the mutation burden across the human genome is highly uneven – captured by the existence of hypo-mutated genes and longevity-associated hypo-mutated pathways that have significantly lower mutation burden than expected. We further find that highly connected network hubs are depleted for mutations, while peripheral, condition-specific programs accumulate higher mutation burden. This organized vulnerability is shown to arise through two independent mechanisms, transcription-coupled repair and selective filtering, and we quantify each mechanism’s relative contribution to mutation accumulation. We then validate these findings under experimental mutagenesis, supporting intrinsic protective mechanisms over tissue-specific confounders. Our findings refine the somatic mutation theory of aging: organismal decline may result not from the overall mutational burden itself, but from the accumulation of mutations specifically within essential genes and network modules, where even rare mutational incursions could have outsized phenotypic consequences.

## Results

### Somatic mutations accumulate non-randomly across genes

We begin by analyzing 1.3 million mutations mapped to 18,547 protein-coding genes from SomaMutDB [34] including intron and exon regions, encompassing 4,684 cells from 183 individuals across 13 tissues (Methods 1). We find that mutation counts correlate strongly with gene length (*r* = 0.93, *p* < 10^−12^; Fig. 1a), prompting us to use a length-normalized Monte Carlo null to isolate deviations that reflect meaningful deviation rather than random chance (see Methods 2). This analysis identified 727 hypo-mutated genes (3.9%; i.e. genes with less mutations than expected) and 1,253 hyper-mutated genes (6.8%; i.e. genes with that exceed expected number of mutations) after multiple testing correction (Fig. 1b; FDR *⍺* = 0.05, | log_2_ FC| ≥ 1.5). For example, *IFNL2*, encoding an interferon involved in antiviral immune signaling at epithelial barriers [35], is hyper mutated, accumulating nearly 31-fold more mutations than predicted by length alone (log_2_ FC = 4.95, *p* < 1 × 10^−5^). By contrast *USP9Y*, a deubiquitinating enzyme, is hypo-mutated, with only 2.1% of its expected mutations (log_2_ FC = −5.54, *p* < 1 × 10^−5^) .

**Figure 1:**
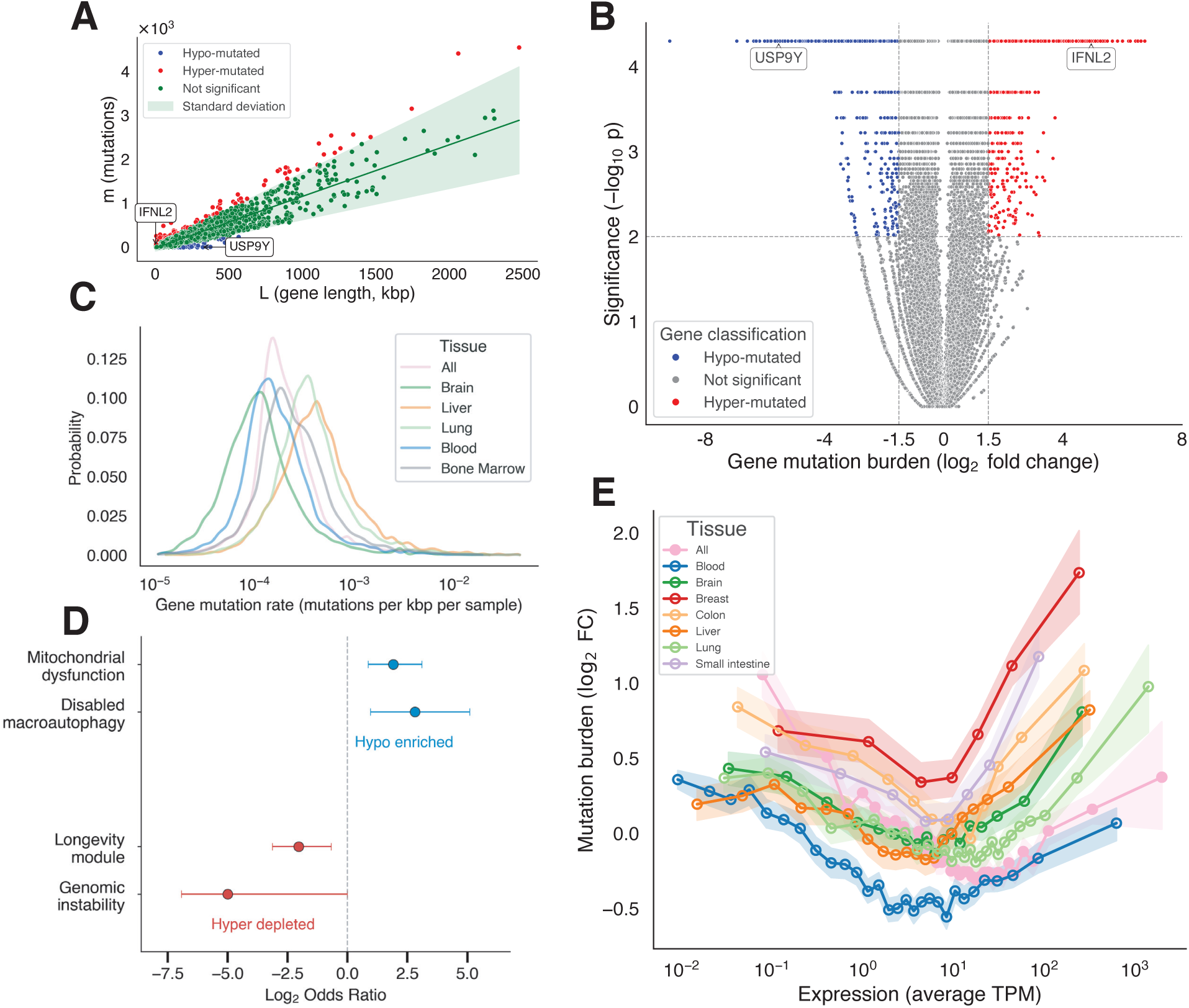
Mutational burden across genes. (**A**) We find a strong linear correlation (*p* = 0.94, *p* < 10^−12^ Pearson correlation) between gene length and number of mutations per gene, where each point is a gene and the green line is the linear model with the standard deviation of the data. (**B**) The 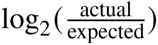 of somatic mutation burden (fold change or FC) vs. the empirical *p* value to identify the h per-mutated (red), hypo-mutated (blue) and not significant (gray) genes according to the length based null model. (**C**) The distribution of mutation rate (mutations/kbp/sample) of genes across different tissues each with at least 100 unique samples. (**D**) Enrichment of hallmarks of aging associated genes with hyper- and hypo-mutated genes. Mitochondrial dysfunction and disabled macroautophagy are enriched with hypo-mutated genes, whereas the full longevity gene set (all genes associated with aging) and genomic instability are significantly depleted in hyper-mutated genes, both implying an overall lack of mutations. (**E**) Somatic mutation burden (log_2_ FC as a function of expression measured in TPM across tissues with expression data. Each point represents the average expression and mutation burden for an expression-based bin of 500 genes with the standard error shaded. Despite an overall negative correlation (*r*_*Spearman*_ = −0.21, *p* = 1.09 × 10^−175^), both low and highly expressed genes show elevated burden, forming a U-shaped pattern across all tissues.

### Gene-specific mutation burden is consistent across tissues

Our data allows us to confirm that tissue-specific mutation rates span more than an order of magnitude, from 1.8 mutations per year in heart and 2.4 per year in brain to 11.5 per year in liver and 10.3 per year in small intestine (Fig. 1c, Fig. 1a) [23]. These patterns raise the question of whether hypo-mutation occurs in a tissue-independent manner, regardless of mutation rate. Consequently, we identified 77 consistently hypo-mutated and 67 consistently hyper-mutated genes, representing 71-fold and 25-fold enrichments over random expectation, respectively (both *p* < 0.001; see Methods 2; Supplementary Data S1; Fig. 1b). Consistently hypo-mutated genes (e.g., *PRH1*, *GCOM1*) are enriched for cell adhesion and developmental processes, while consistently hyper-mutated genes (e.g., *KRTAP19-1*, *OR14I1*) are enriched for sensory perception and G-protein-coupled receptor activity, the latter reflecting peripheral, condition-specific functions (see Supplementary Text Sect. 1)

Critically, 44 genes (0.24%) exhibited mixed mutational behavior across tissues, 12.5-fold fewer than the 572 expected by chance (*p* < 0.001). This depletion demonstrates that genes are systematically predisposed toward or against mutation accumulation independent of tissue context, reflecting intrinsic differences in DNA repair allocation or selective constraint rather than tissue microenvironment alone.

### Longevity genes show non-random mutational patterns

Somatic mutation accumulation has been linked to aging and lifespan, prompting us to hypothesize that longevity-associated genes would exhibit non-random mutational patterns. We assembled a longevity gene set consisting of 286 genes manually curated by Open Genes [36], that allows us to group genes linked to the 11 established hallmarks of aging, and to assess over-enrichment using Fisher’s exact tests [3, 36, 37] (see Methods 2). We find that hypo-mutated genes are strongly enriched in Mitochondrial dysfunction (OR=3.75, *p* = 0.0045) and Disabled macroautophagy (OR=7.00, *p* = 0.046; Fig. 1d). Conversely, hyper-mutated genes are significantly depleted from the longevity gene pool (OR=0.24, *p* = 1.78 × 10^−4^) as well as for gene sets associated with hallmarks of aging, like Genomic instability (OR=0.0, *p* = 0.048) and Epigentic alterations (OR = 0.30, *p* = 0.093) achieving marginal significance. The selectivity of these signals is itself biologically informative. Mitochondrial function and macroautophagy, the two hallmarks that show protection against mutation, share two properties that directly engage protective mechanisms: constitutive, broad expression across cell types (driving transcription-coupled repair) and immediate, cell-autonomous fitness consequences upon disruption (driving purifying selection) (Supplementary Text Sect. 2). This suggests that organized protection concentrates on cellular infrastructure where a single mutational event has no compensatory alternative.

### Non-linear relationship between transcription and mutation burden

While gene expression is known to modulate mutation heterogeneity [38, 39, 40], its direction is affected by competing effects, resulting in conflicting outcomes in prior studies: some studies document negative correlation between expression and normalized somatic mutation burden, attributed to TCR-mediated protection of highly transcribed loci [26, 39], while others see positive correlations, attributed to transcription-associated mutagenesis [41, 42]. To test the net outcome of these opposing mechanisms, we collected median gene expression (TPM) from GTEx across 10 tissues [43], matched to the tissues in SomaMutDB and compared with the normalized mutation burden (see Methods 3). We observe a robust U-shaped nonlinear relationship (Fig. 1e). This pattern was consistent across all tissues, indicating that opposing forces—TCR protection at moderate expression and transcription-induced damage at high expression—act through tissue-independent mechanisms. The U-shape suggests that the previous conflicting conclusions [26, 39, 41, 42] need not be contradictory, but observed a different segment of the expression spectrum (see Supplementary Text Sect. 3). The U-shape reveals a fundamental paradox of cellular maintenance: while moderately expressed genes are protected below baseline, the genes a cell expresses most heavily are simultaneously the best-repaired and the most damaged, as transcriptional machinery that recruits repair also generates the very lesions repair must fix. Beyond a critical expression threshold, the cell’s own activity becomes a source of genomic vulnerability it cannot fully resolve, suggesting that the most transcriptionally demanding genes may mark the breaking points where protection eventually fails with age (see Supplementary Text Sect. 3). On a more technical level, this non-linearity helps us establish expression-matched controls for subsequent pathway analyses.

### Pathways show protection beyond transcription

Since genes are fundamentally linked to biological function through their organization into pathways, next we asked whether such functional modules exhibit coordinated mutation patterns beyond what expression alone predicts. Specifically, for 1,481 Reactome pathways [44], we compare the pathway’s mutation burden against 10,000 surrogate gene sets matched for both size and expression level, ensuring that any signal reflects functional pathway membership rather than shared transcriptional activity (see Methods 4). While 95.7% of pathways showed no deviation from expectation, we identified 39 significantly hypo-mutated pathways after multiple testing correction (Fig. 2a,b), containing much fewer mutations than expected by chance. This pathway-level hypo-mutation is achieved through a segregation pattern: hypo-mutated genes are 1.88-fold enriched within these pathways (*p* = 0.012), while hyper-mutated genes are absent (*p* = 3.1 × 10^−9^).

**Figure 2:**
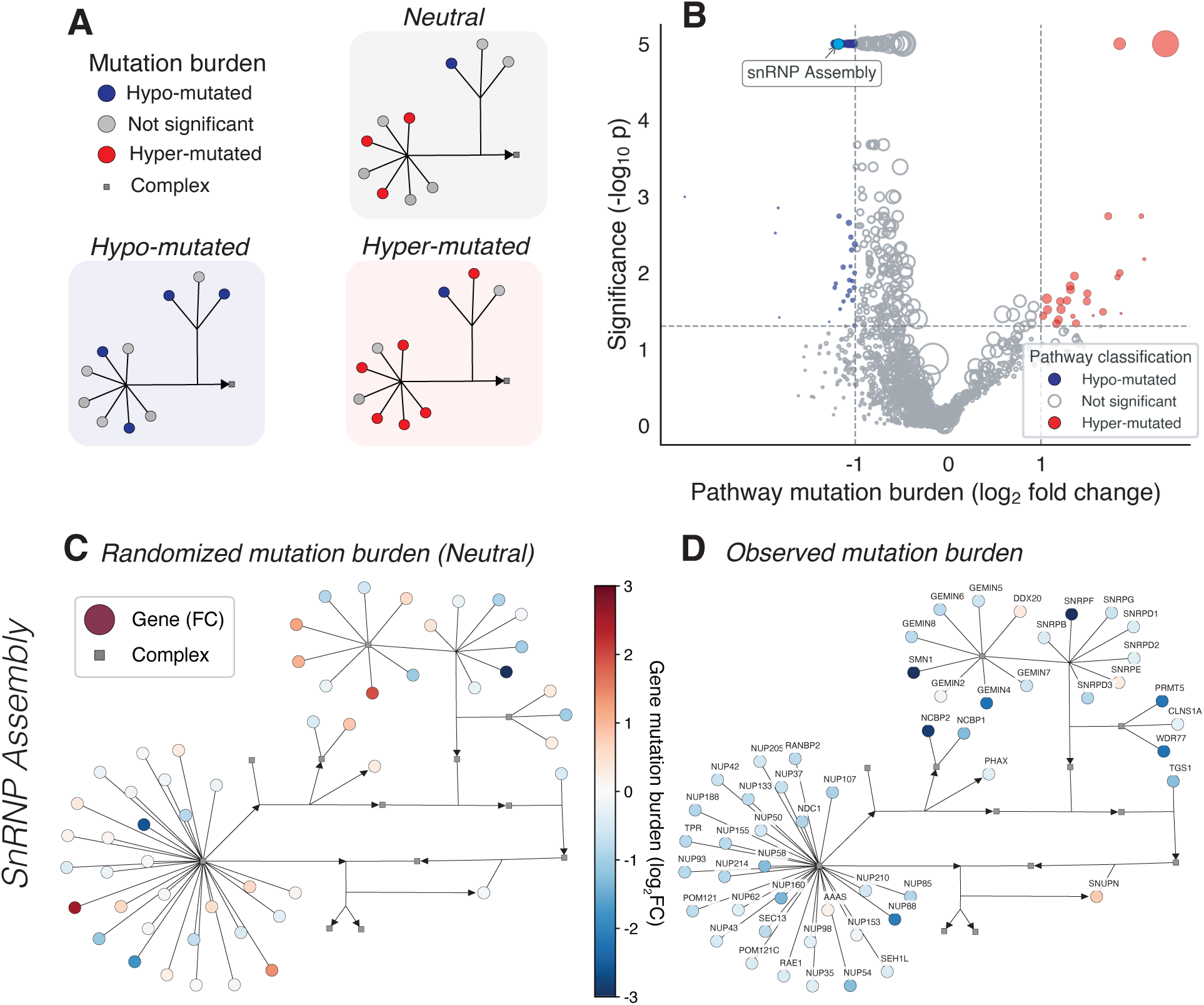
Hyper- and hypo-mutated pathways. (**A**) The classification of pathways based on their mutational burden into neutral (not significant), hypo-mutated, and hyper-mutated. (**B**) Volcano plot of pathway mutation burden, measured as the log_2_ foldchange, vs. the significance, where each point is a pathway. Symbol size is proportional to the number of member genes. While 95.7% of pathways are neutral, 39 are significantly hypo-mutated (blue) and 25 are significantly hyper-mutated (red). (**C**) Transcription-controlled randomized version of the mutation burden in the snRNP assembly pathway, a significantly hypo-mutated pathway essential for RNA splicing. (**D**) Observed mutation burden in the snRNP assembly pathway. Compared to the randomized version, hyper-mutated genes are almost entirely absent and hypo-mutated genes are enriched, demonstrating that protection is a coordinated property of the pathway module rather than the result of individual gene outliers.

The snRNP assembly pathway exemplifies such coordinated protection. This pathway, essential for RNA splicing and recently linked to aging [45] and Alzheimer’s disease [46], was significantly hypo-mutated (*p* < 10^−4^, log_2_(FC) = −1.18; Fig. 2c,d). Under random expectation defined by expression-matched surrogate gene sets, we predicted 2.1 hypo-mutated and 3.6 hyper-mutated genes; instead, we observed 5 hypo-mutated genes and no hyper-mutated genes (*p* = 0.048 and *p* = 0.056, respectively; Fig. 2a).

We also find 25 significantly hyper-mutated pathways, that show 9.90-fold enrichment of hyper-mutated genes (*p* = 2.0 × 10^−132^) and a depletion of hypo-mutated genes (OR=0.60, *p* = 0.033). Hypo-mutated pathways support essential, continuously required processes—including RNA metabolism, cell adhesion, and core biosynthesis—while hyper-mutated pathways govern adaptive or condition-specific functions such as sensory perception and immune signaling (see Supplementary Text Sect. 1; Fig. 2b,c). This dichotomy indicates that the aging genome is not uniformly affected by somatic mutations; rather, mutation burden reflects a balance between maintaining stability in functionally important systems and permitting variability where mutational cost is lower.

### Network hubs accumulate fewer mutations

If cellular network position reflect functional importance, then the mutational protection observed at the gene and pathway level should be mirrored in the network topology. To examine how mutational heterogeneity maps onto the cellular network architecture, we analyzed the position of hyper-and hypo-mutated genes within the human protein-protein interaction (PPI) network [47, 48] (see Methods 5). The 18,168 PPI network, consisting of experimentally-detected physical binding inter-actions, exhibits a well-documented degree-based heterogeneity, with highly connected hubs maintaining global connectivity while peripheral proteins contributing to localized functions [49, 50]. Prior work established that hub deletion is disproportionately lethal in yeast [51], hubs accumulate fewer amino-acid-altering substitutions over evolutionary time [52], are preferentially protected by redundant gene duplicates [53], and tend to be more broadly and constitutively expressed than peripheral proteins [54]. Whether this functional privileging of network hubs is reflected in somatic mutation patterns during normal human aging is unknown. Here, we discover a clear negative relationship between node degree and mutation burden (*r*_*Spearman*_ = −0.195, *p* = 5.9 × 10^−150^; Fig. 3a): highly connected proteins are systematically depleted of mutations, whereas peripheral proteins accumulate excess mutations. In other words, network centrality serves as a proxy for functional indispensability: mutations in highly connected proteins risk cascading failures across multiple cellular processes, hence are depleted; whereas peripheral proteins, operating in more isolated functional contexts, tolerate greater mutational load without systemic consequence.

**Figure 3:**
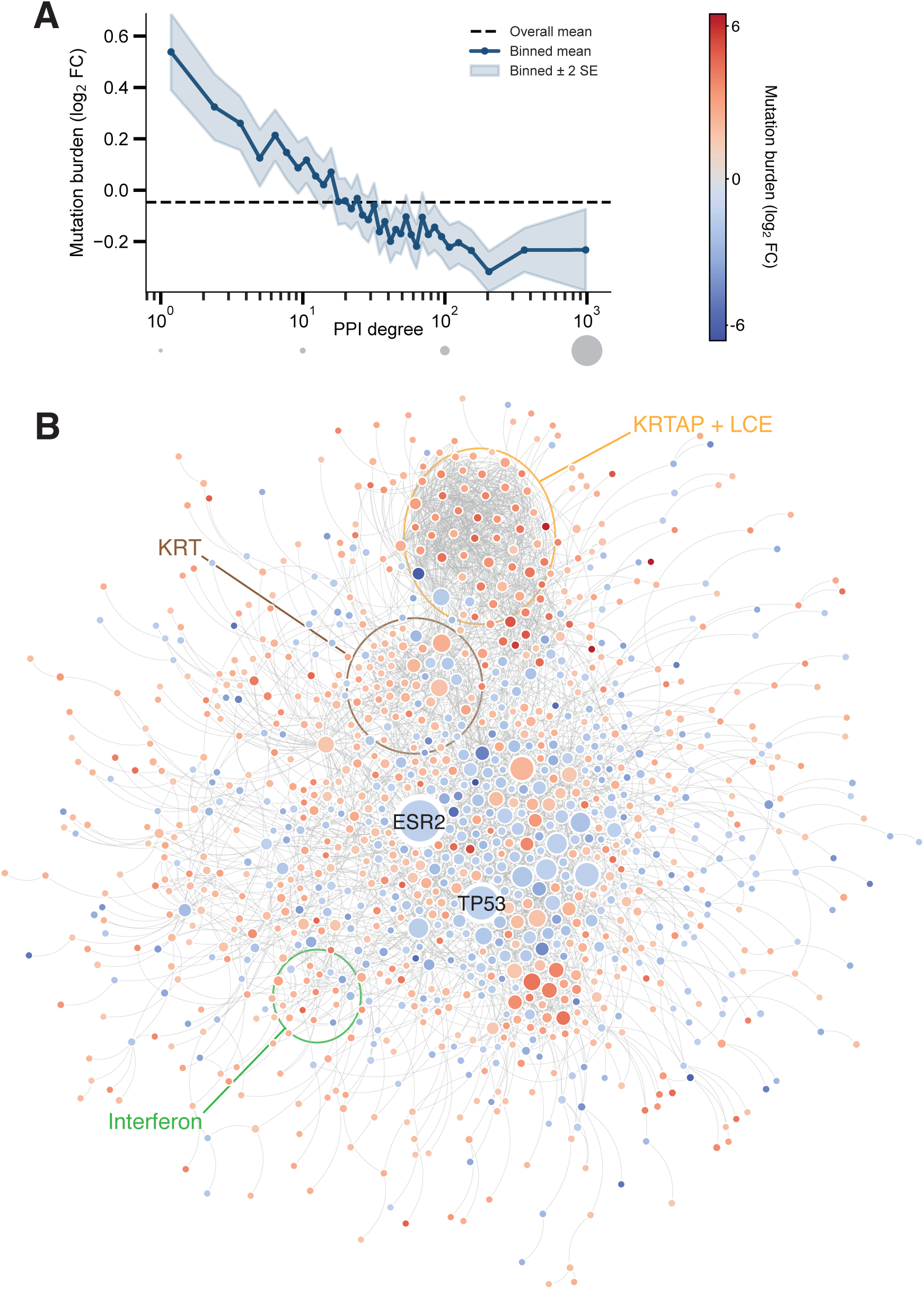
Network organization of hyper- and hypo-mutated genes. (**A**) Mean mutation burden (log_2_(FC)) as a function of node degree in the protein–protein interaction (PPI) network (*r*_*Spearman*_ = −0.195, *p* = 5.9 × 10^−150^). Peripheral proteins (degree 1–10) carry 22% elevated mutation burden relative to expectation, while network hubs (degree ≥ 100) show 13% depletion, demonstrating that functional indispensability is systematically reflected in somatic mutation patterns during aging. Shaded regions indicate ±2 standard errors within degree bins. (**B**) Network view of mutation burden across the human Interactome. Node color encodes mutation burden (log_2_ FC); node size scales with the degree. We find that hyper-mutated genes cluster at the network periphery and form a statistically significant largest connected component (LCC; 492/1,024 genes; *p* = 0.014), enriched for keratin-associated proteins (KRTAP, LCE), canonical keratins (KRT), and interferon-related proteins representing specialized, condition-specific functions. By contrast, hypo-mutated genes, such as *ESR2* and *TP53*, are dispersed throughout the high-degree core (average degree 59.43 vs. 36.81 for hyper-mutated; *p* = 0.032), occupying topologically central positions whose perturbation can propagate broadly across cellular processes.

We also find that hyper-mutated genes form an interconnected module located mainly at the network periphery [55]. Testing interaction densities using degree-preserving null models, we found that hyper-mutated genes exhibit 2.08-fold enrichment for interactions with other hyper-mutated genes (*p* < 10^−4^), forming a degree-corrected statistically significant largest connected component (LCC, 492/1,024 genes; *p* = 0.014; Fig. 3b, Fig.3a). This LCC contains densely connected submodules enriched for keratins (KRTAP, LCE, KRT) and interferon-related proteins, reflecting specialized tissue- or state-specific processes. In contrast, hypo-mutated genes, such as *ESR2* and *TP53*, are dispersed throughout the high-degree network core (average degree=59.43 for hypo-mutated vs. 36.81 for hyper-mutated; *p* = 0.0316), occupying topologically central positions that allow perturbations to propagate broadly and lack a statistically significant LCC (Fig. 3b; Supplementary Text Sect. 4).

### Purifying selection drives mutation depletion in essential genes

The observed mutation burden reflects the cumulative outcome of two core processes: the first is DNA damage and repair, which shape the initial mutational landscape, and the second is selective filtering, which depletes cells carrying deleterious mutations through apoptosis, senescence, or competitive disadvantage. This prompts us to test the degree to which selective pressure contributes to the uneven mutation patterns we observed. For this, we quantified gene-specific selection using *dN*/*dS* ratios from dndscv [56, 57], which compare nonsynonymous (amino-acid-altering) to synonymous mutation rates (see Methods 6). We estimated log_2_(*dN*/*dS*) for 9,826 genes with sufficient coding-sequence coverage: values below 0 indicate negative (purifying) selection, where disruptive mutations are actively purged; values above 0 indicate positive selection, where nonsynonymous changes are retained or expand.

We find that missense *dN*/*dS* correlates strongly with normalized mutation burden (*r*_*Spearman*_ = 0.262, *p* = 1.53 × 10^−154^; Fig. 4a): genes with low *dN*/*dS* (strong purifying selection) were systematically hypo-mutated, whereas genes with high *dN*/*dS* show elevated mutation burdens. Specifically, hypo-mutated genes exhibit median *dN*/*dS* of 1.25, compared to 2.40 for hyper-mutated genes (*p* = 1.87 × 10^−25^, Mann-Whitney U) and 1.37 for neutral genes (*p* = 0.25, Mann-Whitney U). While this correlation could partially reflect reduced mutation opportunity in protected genes, the magnitude and consistency across gene classes indicate that selective filtering substantially shapes the observed mutational landscape.

**Figure 4:**
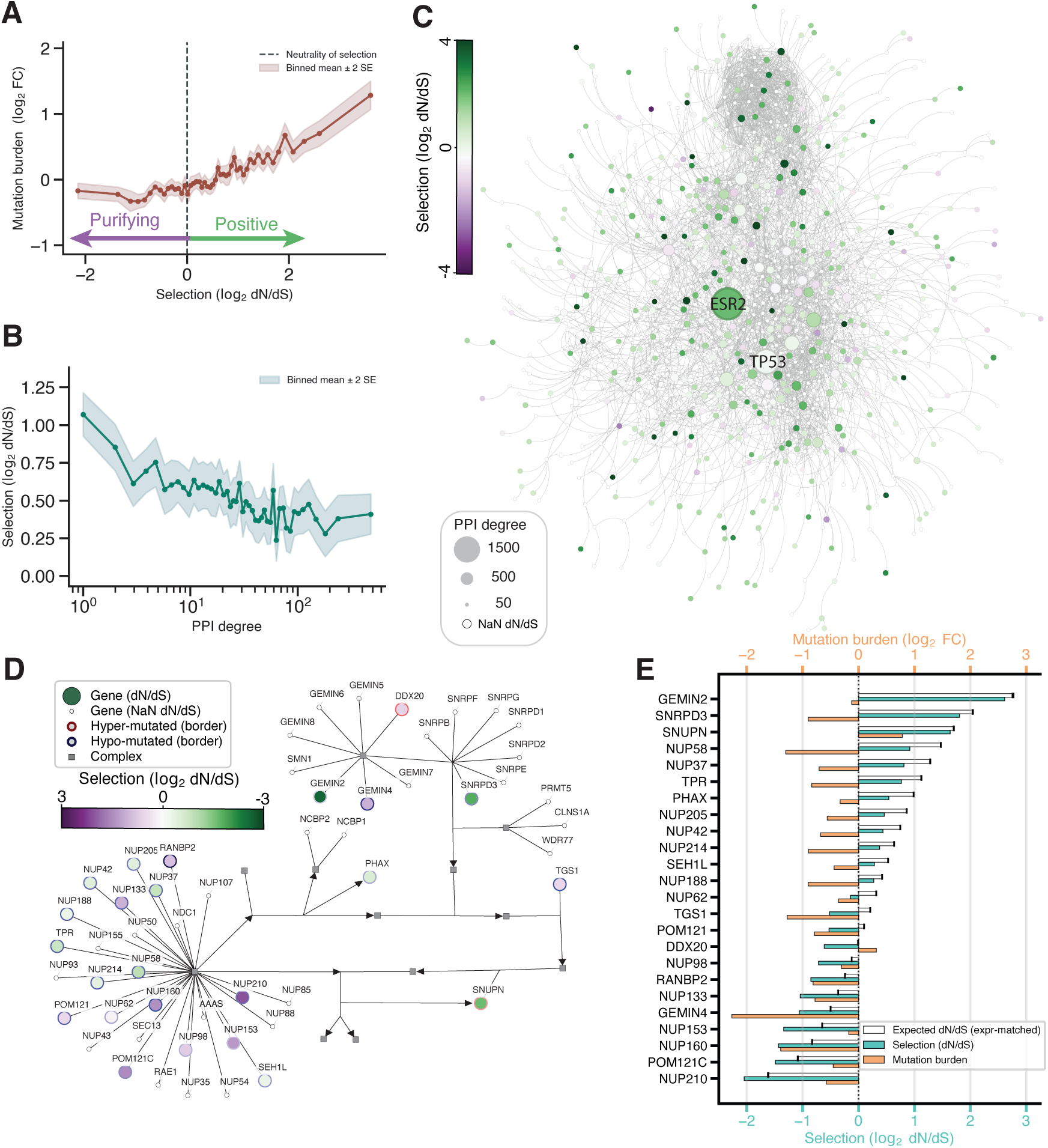
Selective pressures mirror somatic mutation burden and network topology. (**A**) Relationship between missense *dN*/*dS* (log_2_ scale) and somatic mutation fold change across 9,826 genes. Genes under strong purifying selection (low *dN*/*dS*) are systematically hypo-mutated, while genes under weak constraint accumulate excess mutations (*r*_*Spearman*_ = 0.262, *p* = 1.53 × 10^−154^). (**B**) Missense *dN*/*dS* decreases with protein degree in the protein (*r*_*Spearman*_ = −0.119, *p* = 4.68 × 10^−31^), showing that network hubs experience stronger purifying selection than peripheral proteins. (**C**) Interactome colored by missense *dN*/*dS* value, with sizing proportional to Interactome degree. White nodes indicate genes with undeterminable *dN*/*dS*. (**D**) snRNP assembly pathway colored by missense *dN*/*dS*, with borders colored by mutation burden. Most genes exhibit low *dN*/*dS*, reflecting strong purifying selection coordinated at the module level rather than driven by individual outliers. (**E**) For nearly all snRNP assembly genes, observed *dN*/*dS* values fall below expression-matched random expectation, confirming that coordinated evolutionary constraint is an intrinsic property of this pathway and not an artifact of its expression level or network position.

Selective pressure also aligned with network topology. Among 9,426 genes with both PPI and *dN*/*dS* data, node degree correlated negatively with *dN*/*dS* (*r*_*Spearman*_ = −0.119, *p* = 4.68 × 10^−31^; Fig. 4b,c), indicating that network hubs show weaker positive selection (median *dN*/*dS* = 1.25 for degree ≥ 100; 1,360 genes) compared to peripheral proteins (median *dN*/*dS* = 1.59 for degree 1-10; 2,046 genes; *p* = 2.17 × 10^−17^). This mechanistically explains why hubs accumulate fewer mutations—they experience both enhanced repair (via higher transcription) and stronger negative selection against protein-altering changes.

At the pathway level, the snRNP assembly pathway helps to illustrate these coordinated selective constraints. Compared to size- and expression-matched random gene sets, snRNP assembly genes displayed consistently lower *dN*/*dS* values than the random expectation (*p* = 0.034; Fig. 4d,e), demonstrating that purifying selection operates as a collective property across functionally coherent modules rather than being driven by individual outliers.

### Expression and selection independently determine mutation burden

Expression, network degree, and selection each correlate with mutation burden, but these variables are themselves intercorrelated. To unveil their relative contributions, we fit generalized linear models to predict gene-level mutation burden from (i) expression (log_10_ TPM), (ii) selection (log_2_ *dN*/*dS*), and (iii) network connectivity (log_10_ degree), restricted to 9,134 genes with all three annotations available (see Methods 7).

Single-predictor models reveal that expression explains the largest fraction of variance (*R*^2^ = 0.160), substantially exceeding both *dN*/*dS* (*R*^2^ = 0.069) and degree (*R*^2^ = 0.052; Fig. 5a). Network hubs and genes under strong purifying selection might be expected to capture redundant sources of protection, as both mark functionally indispensable genes by independent criteria, topology and evolution. Pairwise models, however, reveal the opposite: expression and *dN*/*dS* together provide largely independent information as their combination yielded the best two-variable model (*R*^2^ = 0.201), while degree contributes minimally beyond expression alone (*R*^2^ = 0.165). This indicates that degree’s association with mutation burden operates primarily through shared variance with expression, while selection contributes an orthogonal component.

**Figure 5:**
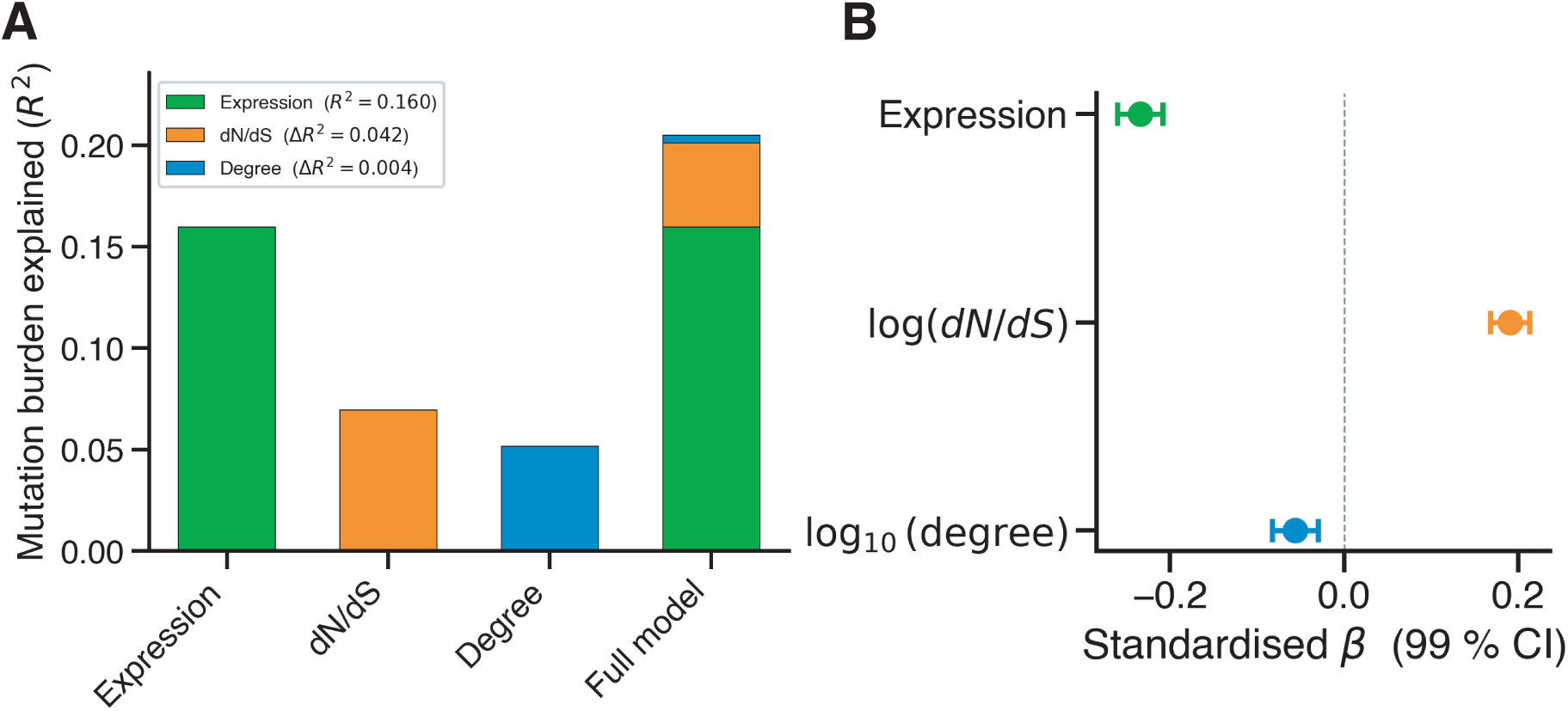
Predicting the origin of mutation burden. (**A**) Variance in gene-level mutation burden explained (*R*^2^) by single-predictor and combined models incorporating expression (log_10_), selective constraint (log_2_ *dN*/*dS*), and network connectivity (log_10_ degree). We find that expression dominates (*R*^2^ = 0.160), selection adds substantial orthogonal variance (Δ*R*^2^ = 0.042), and degree contributes negligibly once the other two are included (Δ*R*^2^ = 0.004), consistent with hub protection operating through expression-dependent mechanisms. The full model explains *R*^2^ = 0.205. (**B**) Standardized coefficients (*β*, 99% CI) from the full model. All three predictors are statistically significant, but their effect sizes reflect the variance decomposition: expression (*p* = 1.42 × 10^−114^) and selection (*p* = 2.17 × 10^−102^) carry the dominant signal while degree’s contribution (*p* = 3.50 × 10^−8^), though highly significant, is substantially smaller.

In the full model incorporating all three predictors (*R*^2^ = 0.205), all three coefficients remained significant (Expression: *β* = −0.2346, *p* = 1.42 × 10^−114^; *dN*/*dS*: *β* = 0.1904, *p* = 2.17 × 10^−102^; Degree: *β* = −0.0562, *p* = 3.50 × 10^−8^; Fig. 5b). However, their incremental contributions differed markedly: expression explained Δ*R*^2^ = 0.160, *dN*/*dS* explained Δ*R*^2^ = 0.042, and degree added only Δ*R*^2^ = 0.004 beyond the other two predictors (see Supplementary Text Sect. 5).

Together, these factors explain ∼20% of gene-level variance in mutation burden, suggesting that additional mechanisms, including chromatin accessibility, replication timing, local sequence context, and stochastic processes, contribute substantially to the mutational landscape of aging.

### Experimental mutagenesis validates protective mechanisms

If hypo-mutation reflects intrinsic gene-level protection rather than tissue-specific context, these patterns should persist under experimentally induced mutagenesis. To test whether mutation patterns reflect stable gene-intrinsic properties, we induced somatic mutations in primary human fetal lung fibroblasts (IMR-90) through repeated sublethal exposure to N-ethyl-N-nitrosourea (ENU; 9 cycles, 50 µg/mL; see Methods 8) [28]. A key confound in observational data is uneven sequencing coverage, where genes sequenced less deeply will appear to carry fewer mutations by default. Unlike the aggregated multi-study *in vivo* dataset, this controlled experiment allows the effective length (callable bases) per gene to be measured precisely, enabling accurate normalization in the null model (see Methods 8). Deep sequencing revealed 104,324 mutations across 11,753 genes, enabling direct comparison with *in vivo* patterns under a controlled, uniform mutagenic challenge.

Gene-level classification identified 248 hyper-mutated (2.2%) and 118 hypo-mutated genes (1.1%) after multiple-testing correction (Fig. 6a), confirming that non-random mutation patterns persist under controlled experimental conditions. Gene mutation fold-changes correlated significantly between the ENU and the *in vivo* datasets (*r*_*Pearson*_ = 0.4139, =*p* < 1 × 10^−300^; Fig. 6b). Hypo-mutated genes showed marginally significant concordance across conditions: 5 genes overlap (*HBG2*, *MRPL33*, *PCDHA2*, *PCDHA5*, and *PCDHGA3*) versus 2.26 expected by chance (2.29-fold enrichment, *p* = 0.076). Hyper-mutated genes showed much more consistent concordance: 54 genes overlapped versus 10.1 expected (7.44-fold enrichment, *p* = 3.35 × 10^−25^), indicating that a substantial and reproducible subset of genomic regions are constitutively mutation-prone across both endogenous aging and acute chemical mutagenesis. GO analysis of shared hyper-mutated genes revealed strong enrichment for olfactory receptor and G protein-coupled receptor signaling activity—peripheral, sensory functions consistent with the *in vivo* pattern of vulnerability concentrated in specialized, condition-specific genes.

**Figure 6:**
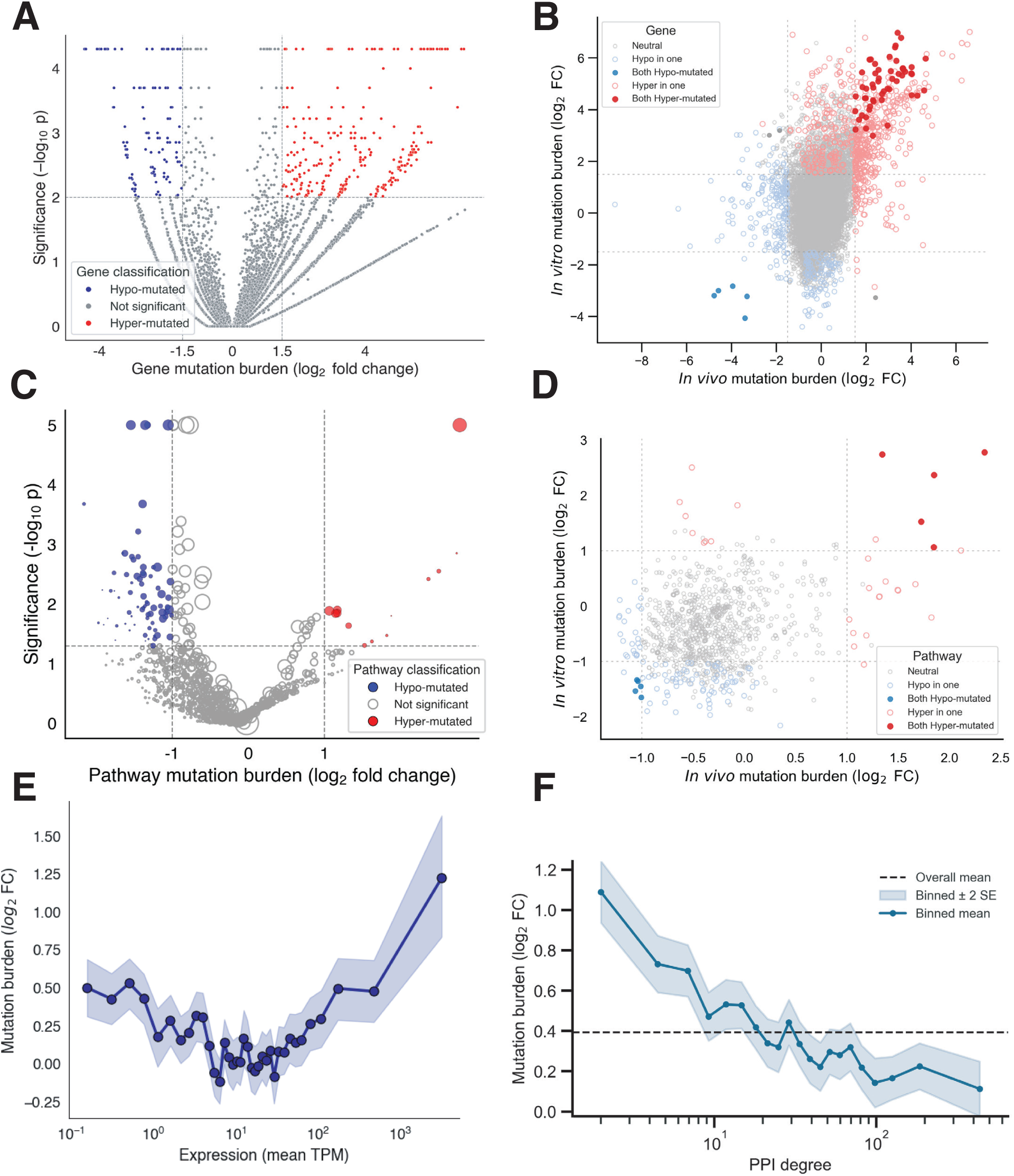
Experimentally mutagenized human samples confirm the *in vivo* mutation patterns. (**A**) Volcano plot showing mutation burden (log_2_ FC) vs. − log_10_(*p*), allowing us to identify hyper-(248, 2.2%) and hypo-mutated genes (118, 1.1%) under controlled ENU mutagenesis of primary human fibroblasts using the same length-based null model as the *in vivo* analysis. (**B**) Correlation between the gene-level mutation burden of ENU-treated cells (vertical axis) vs. the mutation burden observed for *in vivo* aging tissues (horizontal axis; *r*_*Pearson*_ = 0.4139, *p* < 1 × 10^−300^; (see Fig. 1b)). Hyper-mutated genes show strong concordance across conditions (7.44-fold enrichment, *p* = 3.35 × 10^−25^). Hypo-mutated genes show marginally significant concordance (2.29-fold enrichment, *p* = 0.076). (**C**) Volcano plot showing pathway-level mutation burden under ENU mutagenesis compared with the significance of those deviations. Five hypo-mutated pathways overlap with *in vivo* results (3.12-fold enrichment, *p* = 0.037), with 5 hyper-mutated pathways also overlapping (14.79-fold, *p* = 1.06 × 10^−4^; see Fig. 4a). (**D**) Comparison of the mutation burden log_2_ fold change of the ENU-mutagenized samples vs. the *in vivo* samples in pathways (*r*_*Pearson*_ = 0.271, *p* = 4.88 × 10^−18^). (**E**) Expression in bulk TPM versus mutation burden in ENU-treated cells. The nonlinear U-shape is preserved, confirming a nonlinear relationship between transcription and normalized mutation burden remains active under chemical mutagenesis. (**F**) Degree and mutation burden relationship in ENU-treated fibroblasts (*r*_*Spearman*_ = −0.136, *p* = 1.0 × 10^−43^).

At the pathway level, we focused on 988 pathways with at least 5 represented genes, finding 73 hypo-mutated pathways and 13 hyper-mutated pathways in ENU-treated cells (Fig. 6c; see Supplementary Text Sect. 6). Five hypo-mutated pathways overlapped with the *in vivo* analysis (3.12-fold over expectation, *p* = 0.037), suggesting that coordinated protection of functional modules may be reproducible across mutagenic contexts. These pathways span DNA repair and chromosomal maintenance and growth factor signaling. Similarly, 5 hyper-mutated pathways overlapped across conditions (14.79-fold over expectation, *p* = 1.06 × 10^−4^), and these pathways, spanning olfactory receptor expression, cornified envelope formation, metal ion buffering, and interferon signaling, are predominantly tissue-specific or stress-response programs, confirming again that vulnerability reflects convergent and reproducible exposure sensitivity. Pathway fold-changes showed a significant positive correlation across conditions (*r*_*Pearson*_ = 0.271, *p* = 4.88 × 10^−18^; Fig. 6d), indicating that the overall landscape of pathway-level protection and vulnerability is broadly conserved between somatic aging and exogenous mutagenesis, with deviations reflecting differences in damage intensity rather than categorical shifts in which modules are protected.

Bulk expression levels measured within the ENU-mutagenized samples recapitulate the U-shaped relationship observed *in vivo* (Fig. 6e), confirming that both TCR-mediated protection at moderate expression levels and overwhelming transcription-induced damage at high expression levels persist under experimental mutagenesis. The consistency of this pattern across observational and controlled settings indicates that the opposing forces underlying the U-shape operate through intrinsic, damage-independent mechanisms.

We also find that network degree continues to correlate significantly with mutation burden in ENU-treated fibroblasts (*r*_*Spearman*_ = −0.136, *p* = 1.0 × 10^−43^; Fig. 6f) and *dN*/*dS* did not correlate with degree (*r*_*Spearman*_ = −0.068, *p* = 0.0745; Fig. 4b). This mild attenuation of the degree-mutation relationship in IMR-90 cells closely resembles the pattern observed *in vivo* lung tissue (Fig. 1d).

Gene-level sparsity in the ENU dataset limits the power of a full selection analysis; nevertheless, we conducted this analysis and report the results in the Supplementary Text (Sect. 6). Taken together, the experimental data recapitulate the key signatures observed *in vivo* – non-random mutation patterns, network-degree-dependent protection, and transcription-coupled repair – confirming that genomic vulnerability during aging reflects stable, intrinsic properties of gene architecture rather than artifacts of observational context.

## Discussion

Since Szilard’s 1959 hypothesis linking somatic mutations to aging, the prevailing assumption has been that DNA damage accumulates stochastically across the genome. Here we demonstrate that the aging genome exhibits organized vulnerability rather than uniform degradation: somatic mutations accumulate non-randomly, so that essential genes, longevity-associated pathways, and highly connected network hubs are systematically protected, while specialized, adaptive programs tolerate higher mutation burdens.

The protective organization we observe operates through mechanistically distinct pathways. Transcription-coupled repair and related expression-linked processes dominate, accounting for approximately three times more variance than selection. This is consistent with decades of work establishing that actively transcribed genes receive preferential DNA repair [38, 40], but represents the first systematic quantification in normal somatic aging rather than cancer or experimental damage. The U-shaped relationship between expression and mutation burden reveals competing forces: transcription-coupled repair protects moderately expressed genes, while intense transcription generates damage through open chromatin, replication-transcription conflicts, and R-loop formation. Selective filtering contributes independently: genes under strong purifying selection (low *dN*/*dS*) show reduced mutation burden even after accounting for expression, and this relationship strengthens under experimental mutagenesis when environmental noise is eliminated. Critically, network topology’s apparent protective effect operates entirely through expression-dependent mechanisms: highly connected hubs are protected due to their expression levels. This key finding was validated under experimental mutagenesis in a single cell type, where the U-shaped expression-mutation relationship and network-degree-dependent protection were both preserved.

Several features of our results indicate that protection reflects stable, gene-intrinsic properties rather than tissue-specific artifacts. First, the same genes and pathways are systematically hypo- or hyper-mutated across tissues despite order-of-magnitude variation in absolute mutation rates. Second, genes consistently hypo-mutated across tissues exceed random expectation, while mixed-behavior genes are depleted, demonstrating systematic commitment to mutational trajectories. Third, controlled chemical mutagenesis revealed an asymmetric concordance: a marginally significant core of hypo-mutated genes was reproduced across conditions, indicating constitutive vulnerability in peripheral genes, representing a condition-independent feature of the mutational landscape.

These findings carry implications for understanding aging at the cellular and organismal levels. The traditional mutation accumulation theory predicts that aging results from stochastic degradation once mutations exceed a threshold. Our data suggest a more nuanced model: aging may result not from uniform mutation burden but from the eventual failure of protective mechanisms in essential modules, leading to disproportionate functional decline when critical pathways accumulate damage. The systematic protection of longevity-associated genes—particularly those involved in mitochondrial function and autophagy—supports this view and suggests that interventions targeting these specific modules may be more effective than genome-wide mutation suppression. The depletion of hyper-mutated genes from longevity pathways and the rarity of mutations in genes like those involved in snRNP assembly (linked to aging and neurodegeneration) indicate that even rare mutational incursions into protected modules could have outsized phenotypic consequences. This framework also reconciles the observation that individuals with elevated somatic mutation burdens due to DNA polymerase defects do not exhibit premature aging: if mutations accumulate preferentially in peripheral, functionally redundant genes, total burden may be less informative than functional distribution across the cellular network.

Our study has limitations that temper interpretation. First, our analysis is observational and cannot definitively establish causality between mutation patterns and aging phenotypes. Second, SomaMutDB does not provide per-gene coverage files, precluding direct correction for sequencing depth heterogeneity across genes and studies; the controlled ENU experiment addresses this limitation through known sequencing coverage, but residual confounding in the observational analysis cannot be fully excluded. Third, our mutation burden metric aggregates mutations across entire genic regions without distinguishing between intronic and exonic variants; alternative approaches that weight or restrict to coding sequences or expand to non-coding sequences may yield different estimates of normalized mutation burden. Fourth, while we interpret expression as a proxy for transcription-coupled repair, we do not directly measure repair activity; the correlation could partially reflect confounding factors such as chromatin accessibility or replication timing. Fifth, our experimental validation used a single cell type and chemical mutagen; testing whether protection persists across diverse cell types and mutational challenges would strengthen generalizability. Finally, virtually all mutations in SomaMutDB are single-nucleotide variants or small insertions and deletions, with the much less frequent genome structural variants, including copy number variants, retrotranspositions and chromosomal level variants, essentially absent.

## Methods

### 1 SomaMutDB data preprocessing

We assembled a unified somatic mutation dataset from SomaMutDB [34] by aggregating all available single-cell and clonal sequencing studies (downloaded on March 29, 2025). Raw VCF files were merged with SomaMutDB sample metadata to obtain tissue, donor age, sex, and sequencing method. Only samples generated using Clone, MDA, PTA, or META-CS protocols were included, while samples representing multiple pooled cells were removed. Mutations were mapped to genes using GRCh38 Ensembl reference genome with pysam (version 113; downloaded March 4, 2025), retaining only unambiguous gene assignments to protein-coding genes. After filtering, the final dataset comprised 1,323,796 somatic mutations from 4,684 cells derived from 183 individuals across 13 tissues, of which 1,279,401 are SNVs. This processed mutation table formed the basis for all downstream analyses.

### 2 Gene-level Monte Carlo null model

To identify genes with non-random somatic mutation burden, we compared observed mutation counts to a gene-length–normalized Monte Carlo null. Across all samples, we estimated a global mutation rate of 1.004 × 10^−3^ mutations per base pair from 4,684 cells, considering protein-coding regions only. Under the null hypothesis of random mutagenesis, each nucleotide position is treated as an independent Bernoulli trial with probability given by this global rate, implying that a gene’s expected mutation burden is determined solely by its length. Gene lengths were derived from nucleotide counts computed on the GRCh38 Ensembl reference genome (downloaded March 4, 2025) and aggregated by gene symbol.

Using this framework, the Bernoulli model at individual base pairs is statistically equivalent to a multinomial formulation in which the observed total number of mutations is redistributed across genes in proportion to their lengths. We therefore generated stochastic realizations of genome-wide mutation patterns by reallocating mutations, weighted according to gene length, repeating this procedure 10,000 times to construct empirical null distributions for each gene. Sampling was performed using NumPy’s multinomial random generator (version: 1.23). Differential mutation burden was quantified as the log_2_ ratio of observed to expected mutations, and empirical two-sided *p*-values were obtained from the simulated distributions and adjusted for multiple testing. Genes were classified as hyper- or hypo-mutated using combined effect-size and significance thresholds (| log_2_ FC| ≥ 1.5, *p* < 0.01).

To assess tissue specificity, we repeated this analysis independently within each tissue using the original tissue annotations provided by SomaMutDB. For each tissue, we recalculated the global mutation rate across genes with at least one observed mutation and applied the same length-weighted Monte Carlo framework to generate tissue-specific null distributions. Genes were classified as hyper- or hypo-mutated within each tissue using identical effect-size and significance criteria. This enabled direct comparison of mutation burden patterns across tissues while preserving a consistent statistical framework.

To assess whether genes implicated in aging are preferentially protected from or targeted by somatic mutation, we curated longevity-associated gene sets from Open Genes [36], which annotates genes across established aging hallmarks, and we retain only genes supported at confidence level ≤4. This yielded a global longevity module together with hallmark-specific subsets. For each set, we tested enrichment among hyper- and hypo-mutated genes using Fisher’s exact test, with the background defined as all genes evaluated in the Monte Carlo framework. Multiple testing correction was performed across all hallmark (11 gene sets) and mutation-class (two mutation classes) combinations using the Benjamini–Hochberg procedure. This analysis was used to identify aging hallmarks exhibiting systematic deviations from random somatic mutagenesis.

### 3 Gene expression and mutation burden

To relate somatic mutation burden to gene expression, we integrated mutation data with median transcript abundance from GTEx v8. Gene-level expression values (TPM) were averaged across available tissues and matched to protein-coding genes included in the mutation analysis. For each gene, mutation burden was quantified using the length-normalized Monte Carlo log_2_ fold-change described above and merged with expression values by gene symbol. Genes were ranked by expression and grouped into equally sized bins (500 genes per bin), within each tissue and for the aggregated dataset. For each bin, mean mutation burden and standard error were computed to visualize trends across the expression spectrum. Only tissues with sufficient gene coverage to support binning were retained. To quantify the association between gene expression and mutation burden, we computed Spearman rank correlations between expression levels and gene-level mutation burden across all protein-coding genes, both for the aggregated dataset and within individual tissues. Correlations were evaluated using SciPy (version: 1.10), providing a nonparametric measure of monotonic dependence between expression and somatic mutation rate.

### 4 Pathway-level mutation burden analysis

Pathway definitions were obtained from Reactome (Reactome.org). Because Reactome is hierarchically organized, we restricted analysis to leaf pathways (those without child pathways) and considered gene–pathway membership without weighting by functional role. After filtering to pathways containing at least two genes represented in SomaMutDB, this yielded 1,481 pathways spanning 9,264 genes.

For each pathway, mutation burden was defined as the mean observed-to-expected gene-level mutation burden across its member genes. Statistical significance was evaluated using an expression-aware surrogate gene set approach. For each pathway of size *n*, we generated 10,000 surrogate gene sets matched in size, in which each pathway gene was replaced by a randomly selected gene of similar expression level (within ±100 expression ranks), thereby controlling for transcription-associated mutation biases. Surrogate distributions provided empirical null models for pathway mutation burden. Two-sided empirical *p*-values were computed from these distributions and adjusted for multiple testing using the Benjamini–Hochberg procedure. Pathways were classified as hyper- or hypo-mutated using an FDR threshold of 0.05 together with an effect-size cutoff of | log_2_ FC| ≥ 1, where the fold change is given by the ratio of observed over expected.

### 5 Protein–protein interaction network analysis

To examine whether hyper- and hypo-mutated genes exhibit non-random network organization, we analyzed their connectivity within a consolidated human protein–protein interaction (PPI) network, where genes are nodes, and the physical interactions between them are edges, assembled from multiple high-throughput and literature-curated sources from six different kinds of protein-protein interactions: (i) Binary PPIs systematically tested by high-throughput yeast two-hybrid (Y2H) experiments (HI-Union [58], HINT [59], HIPPIE [60]); (ii) Kinase–substrate interactions from both literature-curated low-throughput and high-throughput studies (KinomeNetworkX [61], Human Protein Resource Database [HPRD] [62], PhosphoSitePlus [63]); (iii) Literature-curated PPIs identified by affinity purification–mass spectrometry (AP–MS) and low-throughput experiments (InWeb [64], BioGRID [65], PINA [66], MINT [67], IntAct [67], InnateDB [68], APID [69], DIP [70], LitBM17 [58]); (iv) Structure-derived PPIs supported by three-dimensional protein structures or co-fractionation evidence (Instruct [71], Interactome3D [72], INSIDER [73], CoFrac [74]); (v) Signaling and regulatory interactions curated from literature and high-throughput studies (SignaLink2.0 [75], ENCODE [76], RAIN [77], RISE [78]); and (vi) Protein complexes identified through systematic AP–MS experiments or computational inference (BioPlex2.0 [79], QUBIC [80], lncRNome [81], NPInter [82]). The genes were mapped to their official gene symbols based on the National Center for Biotechnology Information (NCBI) database and only genes present in both SomaMutDB and the PPI network were considered. This results in a network with 18,168 genes (nodes) and 520,325 protein–protein interactions.

Network locality was quantified using the size of the largest connected component (LCC) formed by hyper- or hypo-mutated genes within the interactome. To assess statistical significance, we applied a degree-preserving null model in which each gene set was compared against 1,000 randomized gene sets matched both in size and node degree. For each randomization, genes were sampled to reproduce the degree distribution of the gene set. Empirical null distributions of LCC size were constructed from these randomized sets, and z-scores and two-sided *p*-values were computed by comparing observed LCC sizes to the null expectations.

In addition, we examined how gene-level mutation burden relates to network topology by correlating mutation burden (log_2_ fold change) with node degree. Associations were quantified using Spearman correlation coefficients as implemented in SciPy.

### 6 Estimating selective constraint using *dN*/*dS*

To quantify selective pressures acting on somatic variants, we estimated gene-level *dN*/*dS* ratios using dndscv in R [56]. Mutations were formatted for dndscv and sorted by sample, chromosome, and position. To reduce potential inflation from local clustering artifacts, we excluded immediately adjacent mutations (positions differing by one base) occurring within the same sample and chromosome prior to analysis. *dN*/*dS* inference was performed against the GRCh38 coding reference provided by dndscv, with per-gene and per-sample mutation caps disabled (max muts per gene per sample and max coding muts per sample set to Inf) and covariate adjustment disabled. Selection was quantified using the missense-specific *dN*/*dS* estimate (wmis) reported by dndscv. Analyses were performed both on the full dataset and within individual tissues using SomaMutDB annotations.

### 7 Modeling determinants of gene-level mutation burden

Expression, selection, and network connectivity are each associated with mutation burden, but these predictors are intercorrelated. To quantify their independent contributions, we fit generalized linear models (Gaussian errors; identity link, equivalent to linear regression) predicting gene-level mutation burden (log_2_ fold change) from three covariates: expression (log_10_ TPM, computed as above), selective constraint (log_2_ missense dN/dS; wmis from dndscv), and PPI connectivity (log_10_ degree). The analysis was restricted to genes with all three annotations available.

We fit single-predictor, pairwise, and full three-predictor models to compare explained variance across specifications (reported as *R*^2^). To quantify incremental contributions, we computed Δ*R*^2^ for each predictor by comparing the full model to the corresponding reduced model omitting that predictor, and we report standardized coefficients from z-scored predictors for effect-size comparability. Models were fit in Python using statsmodels (version: 0.13).

### 8 ENU experiment

The experimental protocol and data generation for ENU-mutagenized IMR-90 fibroblasts are described in full in Cutler et al. [28]. Our mutation burden analysis followed the same Monte Carlo null framework as the *in vivo* analysis, with two key differences. First, rather than using annotated gene length as the null model denominator, we used an effective gene length derived from sequencing coverage: for each gene, effective length was estimated as the product of the number of covered bases and mean per-base read depth, providing a direct empirical measure of mutational opportunity that accounts for uneven coverage across genes and removes a confound unavoidable in the observational dataset. This was done to account for the uneven coverage inherent in single-cell data. Second, gene expression values were taken directly from pseudobulk RNA measurements within the ENU-mutagenized fibroblast samples themselves rather than from GTEx, ensuring that the expression–mutation relationship reflects the transcriptional architecture of the experimental system rather than population-averaged tissue expression in normal aging human tissues.

## Author contributions

J.E. and A.L.B. conceived of the analysis; J.E. performed computational analysis; R.C. performed wet-lab experiments; R.C., J.S., B.G., O.L., J.V., and X.D. provided guidance on computational analysis; J.E. and A.L.B. wrote the manuscript; all authors reviewed the manuscript.

## Data and materials availability

Code for reproducing the analysis is deposited on GitHub (https://github.com/jpehlert/somatic-mutations-analysis). The observational somatic mutations data is available from SomaMutDB [34]. See [28] for experimental data availability and access.

