## Supplementary material for "The aging genome exhibits organized vulnerability to somatic mutations": SI

### Supplementary Information for *The aging genome exhibits organized vulnerability to somatic mutations*

### Contents

**1 Supplementary Note: Individual tissue signatures reveal organ-specific patterns 2**

**2 Supplementary Note: Biological interpretation of hallmark-specific mutation patterns 4**

**3 Supplementary Note: The nonlinear relationship between transcription and mutation rate 6**

**4 Supplementary Note: Mechanistic basis of hub protection 9**

**5 Supplementary Note: Multifactorial architecture of somatic mutation accumulation 9**

**6 Supplementary Note: Selection analysis and concordance with independent experimental analysis. 10**

**7 Supplementary Data Captions 17**

### 1 Supplementary Note: Individual tissue signatures reveal organ-specific patterns

The aggregate analysis presented in the main text pools somatic mutations across tissues to maximize statistical power, but this raises the question of whether the identified patterns reflect genuine biology or artifacts of tissue mixing. To address this, we examined mutation burden, gene-level classifications, and pathway enrichments separately within individual tissues, finding that the core results replicate consistently across organs despite substantial differences in baseline mutation rates (Fig. 1a).

**Gene-level hypo- and hyper-mutation across tissues.** Across all tissues examined, the proportion of genes classified as hypo- or hyper-mutated remains stable relative to the aggregate (Fig. 1b). All tissues, even those with less than 100 samples consistently demonstrated the same ratio of hyper- to hypo-mutated genes as the aggregated analysis. This consistency holds despite an order-of-magnitude range in tissue-level mutation rates, indicating that the relative architecture of protected and vulnerable genes is a robust feature of the somatic mutation landscape rather than a consequence of overall mutational burden. See Supplementary Data S1 for tissue-by-tissue specific values.

Applying the pathway framework separately within individual tissues revealed coherent, tissue-specific patterns that mirror each organ's functional priorities, using the the transcription-aware null model (Fig. 1c). Below, we report broad themes for three representative tissues: brain, lung, and blood. See Supplementary Data S2 for each pathway within each tissue.

**Brain.** We identified 58 hyper-mutated and 74 hypo-mutated pathways. The strongest hypo-signals concentrate in neuron-specific modules, including *Neuronal System*, synaptic transmission, neurotransmitter receptor signaling, and ion transport. These remain depleted after controlling for expression, consistent with the high functional cost of perturbing excitability and connectivity in largely post-mitotic neurons. In contrast, the hyper-signals are enriched for core maintenance programs—*DNA repair and replication*, *cell cycle*-associated machinery, and transcription/chromatin remodeling—indicating an excess of mutations in genome maintenance and regulatory modules within brain tissue.

**Lung.** We observed 25 hyper-mutated and 46 hypo-mutated pathways. The dominant hypo-mutated set centers on core biosynthetic capacity: *translation/ribosome biogenesis*, *protein processing and secretory trafficking*, and *mitochondrial energy metabolism*. These protected modules align with the sustained metabolic and secretory demands of pulmonary epithelium. The hyper-mutated set again features *DNA repair and cell cycle* modules together with stress-responsive transcriptional regulators, indicating an excess in genome maintenance and stress pathways in lung.

**Blood.** Blood exhibits 29 hyper-mutated and 67 hypo-mutated pathways. Protected pathways are enriched for *translation/ribosome*, *RNA metabolism*, and *hematopoietic differentiation* programs that underpin continuous lineage production. The hyper-mutated set combines the recurrent maintenance axis—*DNA repair/replication* and *cell cycle*—with blood-specific *immune receptor and cytokine signaling* modules, reflecting an excess of mutations in both genome maintenance and immune signaling within hematopoietic contexts.

Two features are consistent across tissues. First, the identity of the hypo-mutated, protected modules tracks tissue identity: brain safeguards neuronal signaling and ion conductance; lung safeguards translational and energetic capacity; blood safeguards protein synthesis and lineage specification. Second, a common hyper-mutated axis recurs across all three tissues—pathway-level excess in *DNA repair/replication*, *cell cycle*, and associated chromatin/transcription modules. This differs from the aggregate analysis, where these same maintenance modules tended to appear among the protected sets. One way to reconcile these perspectives is to recognize that genome maintenance pathways are not confined to a single organ but instead serve the body as a whole. In the pooled analysis, this system-wide role makes them appear as part of the protected core, akin to how each tissue defends its own defining functions. Yet at the level of individual organs, where other priorities dominate, these same maintenance modules emerge as relatively more exposed. The aggregate thus captures their universal importance to organismal survival, while the tissue-resolved view highlights that they also represent a shared axis of vulnerability across diverse cellular environments.

**Network degree and mutation burden across tissues.** The negative relationship between network degree and somatic mutation burden observed in the aggregate analysis holds consistently across all individual tissues with sufficient sample size (Fig. 1d). Blood and bone marrow show the strongest correlations (Spearman $\rho = -0.20$ for both), while liver and lung are more modest ($\rho = -0.09$ and $\rho = -0.10$, respectively), and brain the weakest ($\rho = -0.06$), yet all are statistically significant. That the direction and significance are preserved across tissues with markedly different mutation rates (Fig. 1a) and cellular contexts reinforces that hub protection is a general property of the network architecture rather than an artifact of any single tissue's biology.

**Evolutionary selection and mutation burden across tissues.** The positive relationship between missense selection pressure and somatic mutation burden is likewise consistent across tissues (Fig. 1e), with all tissues showing significant positive Spearman correlations ranging from 0.25 in bone marrow to 0.44 in liver. The variation in correlation strength across tissues likely reflects differences in the intensity of tissue-specific selection constraints and cell turnover rates rather than any breakdown of the underlying mechanism. Together with the degree results, this confirms that the orthogonal relationships identified in the aggregate are not statistical artifacts of pooling heterogeneous tissues, but instead reflect mechanisms operating consistently within and across individual tissue contexts.

### 2 Supplementary Note: Biological interpretation of hallmark-specific mutation patterns

Of the eleven canonical hallmarks of aging, only a subset showed statistically significant enrichment or depletion of hypo- or hyper-mutated genes. This selectivity might initially appear as a limitation—one might expect that all aging-associated modules would be systematically protected if the genome broadly preserves longevity-relevant function. However, we argue that the pattern of which hallmarks show signal, and the direction of that signal, is biologically coherent and mechanistically interpretable in terms of the two protective mechanisms identified in this study: transcription-coupled repair (TCR) and purifying selection.

**Mitochondrial dysfunction genes are hypo-mutated.** Genes in this module are among the most constitutively and broadly expressed in the human genome, as oxidative phosphorylation and mitochondrial quality control are non-negotiable requirements in virtually every differentiated cell type. High, constitutive expression across tissues directly engages TCR machinery, providing continuous preferential repair to this locus class. Simultaneously, the fitness consequences of mutations in these genes are immediate and cell-autonomous: impaired ATP synthesis and dysregulated reactive oxygen species production simultaneously compromise energetic and redox homeostasis, creating strong pressure for apoptotic elimination or competitive disadvantage against unmutated neighbors. Both protective mechanisms therefore converge on this module, consistent with its robust hypo-mutation signal.

**Macroautophagy genes are hypo-mutated.** Autophagy occupies a unique position among aging hallmarks: it is not merely a stress response but a constitutive housekeeping function responsible for clearing damaged proteins, lipid droplets, and organelles—including mitochondria with damaged membranes. Critically, autophagy is also one of the primary mechanisms by which cells clear the downstream consequences of somatic mutation itself (misfolded or dysfunctional proteins). A mutation disabling an autophagy gene therefore creates a feedback catastrophe: the cell loses its capacity to compensate for subsequent damage, dramatically amplifying the phenotypic cost of that single event. This self-referential vulnerability—where the protection machinery must itself be protected—would manifest as exceptionally strong purifying selection, consistent with the high odds ratios we observe. That autophagy genes are both constitutively active (engaging TCR) and subject to extreme fitness penalties upon disruption (engaging selection) explains why this module emerges clearly despite its small gene count (n=9).

**Genomic instability genes are depleted of hyper-mutated genes.** The depletion of hyper-mutated genes from this hallmark (OR=0.0) is conceptually distinct from the enrichment patterns above. DNA repair and genome maintenance genes are not merely essential—a mutation in this class has a multiplicative, genome-wide effect, converting the cell into a mutator phenotype that accumulates damage at every other locus. The selection coefficient against such mutations is therefore among the highest possible, explaining the complete absence of hyper-mutated genes in this module. This is consistent with the well-established observation that biallelic inactivation of mismatch repair genes produces hypermutator phenotypes associated with Lynch syndrome and microsatellite instability cancers—confirming that these genes are under extreme negative selection in normal tissues.

**Why most hallmarks do not show a signal.** The approximately eight hallmarks that show no statistically significant enrichment in either direction share biological features that would reduce the efficacy of both protective mechanisms. Several involve genes that are expressed conditionally or in restricted cell types: senescence effectors are induced by stress rather than constitutively active, telomerase components are actively suppressed in most somatic tissues removing expression-driven protection, and stem cell renewal genes are by definition expressed in a small minority of cells underrepresented in somatic mutation databases. Others involve processes with substantial functional redundancy—intercellular communication pathways, for instance, typically involve large gene families with overlapping ligand-receptor specificities, reducing the fitness cost of any single mutation. Epigenetic regulators occupy an intermediate position: while broadly expressed, many operate within large multi-subunit complexes where partial loss of function is buffered by remaining complex members, attenuating selection pressure on individual components.

This pattern reveals an important principle: the genome does not uniformly protect all biology associated with aging. Instead, protection concentrates specifically where constitutive expression and irreplaceable function coincide. The modules that show the strongest protection are those where TCR and selection act synergistically—broadly expressed enough to receive continuous repair, and functionally isolated enough that no redundant pathway can compensate for their loss. The silence of other hallmarks is therefore not a null result but a mechanistic prediction: organized vulnerability follows the functional architecture of cellular indispensability, not merely the categorical annotation of aging relevance.

### 3 Supplementary Note: The nonlinear relationship between transcription and mutation rate

The relationship between gene expression and mutation rate has been studied across multiple systems, yet the direction of the effect remains contested. Early comparative genomic work in yeast and human germline sequences documented a positive correlation between expression level and mutation rate, interpreted as evidence that transcription-associated mutagenesis (TAM) dominates when integrated over the full expression range [1]. By contrast, analyses of somatic mutations in normal human tissues have more often reported the opposite: highly expressed genes tend to accumulate fewer somatic mutations, an effect typically attributed to TCR-mediated surveillance at transcribed loci [2, 3]. Adding further complexity, cell-type-resolved studies of postmitotic neurons find that somatic mutations are enriched in transcriptionally active genomic regions and exhibit pronounced transcriptional strand asymmetry, suggesting that transcription simultaneously generates lesions and directs their asymmetric repair [4]. Together, this body of work indicates that both processes are real, and that the net sign of the expression–mutation relationship likely depends on the expression range examined, the cell type, and the specific mutational processes under consideration. Our data do not resolve this debate, but they reveal that a global linear summary is itself an oversimplification: the U-shaped relationship we observe suggests that TCR-mediated protection dominates at intermediate expression while transcription-induced damage accumulates at the high end, and that this non-linearity may partially explain why prior studies have reached contradictory conclusions depending on which segment of the expression distribution their data most heavily sampled.

The U-shaped relationship between gene expression and somatic mutation burden implies that transcription may exert qualitatively distinct effects on genome integrity depending on its intensity, with a saturation of TCR occurring [5]. Here we outline one parsimonious mechanistic framework consistent with this observation, which we offer as a speculative hypothesis rather than a proven explanation.

**Low-expression regime: sparse repair.** At the low end of the expression spectrum, mutation rates are elevated. Lowly expressed genes are transcribed infrequently, which may limit the cell's opportunity to deploy transcription-coupled repair (TCR). Because TCR is initiated by stalled RNA Polymerase II (RNAPII), genes that are rarely transcribed likely receive TCR-mediated surveillance only sporadically [6, 7]. Damage arising in these loci—whether from spontaneous hydrolysis, oxidation, or other sources—would then rely on slower, less efficient global genome nucleotide excision repair (GG-NER) or remain unrepaired until replication, plausibly increasing the probability that lesions are converted to fixed mutations.

**Intermediate-expression regime: efficient TCR.** As transcription frequency increases into an intermediate range, mutation rates fall. One consistent explanation is that in this regime, elongating RNAPII more regularly scans the transcribed strand and stalls upon encountering a DNA lesion, signaling damage presence. Such stalling could trigger recruitment of TCR factors (e.g., CSB/ERCC6, CSA/ERCC8) and downstream nucleotide excision repair machinery, enabling more efficient lesion removal before replication [8, 9, 10, 5, 11]. In this window, transcription may act primarily as a surveillance system, consistent with the observed drop in mutation burden.

**High-expression regime: transcription-induced damage overwhelms repair.** Beyond some expression threshold, mutation rates rise again despite presumably continued TCR activity. One possibility is that at very high transcription rates, the rate of transcription-induced damage begins to outpace TCR capacity, though we cannot directly measure this from our data. Several mechanisms from the literature are at least consistent with this idea. Replication–transcription conflicts may become more frequent when RNAPII occupancy is high, potentially leading to fork stalling and double-strand break formation [12]. Persistent R-loops—RNA:DNA hybrids that form co-transcriptionally—could expose the non-template strand to damage and impede replication fork progression [13]. Additionally, topoisomerase I resolves torsional stress generated ahead of RNAPII by introducing transient nicks; at high transcriptional density, these nicks might accumulate or be converted to strand breaks and indels [7]. Whether these mechanisms individually or collectively account for the observed upturn remains unclear.

Taken together, this mechanistic literature appears broadly consistent with the U-shape we observe, but we emphasize that our data do not allow us to distinguish between these possibilities or rule out alternative explanations. For instance, highly expressed genes may simply have distinct chromatin environments, replication timing, or sequence composition that independently elevates their mutation rates. It is also possible that the inflection point varies across tissues or cell types in ways that reflect differences in TCR throughput, R-loop suppression, or replication scheduling—though testing this would require data and experimental perturbations beyond the scope of the present work. We present this framework primarily to illustrate that the observed non-linearity is not obviously inconsistent with known biology, and to suggest that the interplay between transcriptional activity and repair capacity may be a productive direction for future investigation.

### 4 Supplementary Note: Mechanistic basis of hub protection

The consistency of the protected core and the vulnerable periphery directly resolves a long-standing debate in network biology. Batada et al. [14] first raised the possibility that the apparent essentiality and slow evolution of hubs might be proxies for their high expression levels rather than independent consequences of network position – a question that could not be resolved from evolutionary rate data alone. Our experimental design provides a definitive test: in a single cell type under uniform mutagenic challenge, where the expression architecture is fixed, network degree still carries independent protective signal while selection strengthens, confirming that hub protection *in vivo* is entirely mediated through expression-dependent mechanisms. This extends the centrality-lethality principle [15] from gene deletion lethality to somatic mutation accumulation in aging human tissues, while mechanistically grounding it in transcription-coupled repair rather than topology per se.

### 5 Supplementary Note: Multifactorial architecture of somatic mutation accumulation

The limited variance explained by our models ($R^2$=0.20) is itself informative. The 80% unexplained variance indicates that somatic mutation landscapes are shaped by multiple factors beyond expression, selection, and network position. Likely contributors include chromatin accessibility, replication timing, local sequence context, regional variation in repair enzyme activity, and stochastic processes inherent to DNA damage and repair. Some of this variance may also reflect technical noise in mutation detection and expression quantification across heterogeneous datasets. Importantly, $R^2$=0.20 is comparable to variance explained by genetic and environmental factors in many complex biological traits, and the fact that two mechanisms—repair and selection—independently contribute orthogonal variance components suggests a multifactorial architecture. The unexplained variance represents not a limitation of our approach but an opportunity.

### 6 Supplementary Note: Selection analysis and concordance with independent experimental analysis.

Gene-level sparsity in the ENU dataset renders reliable $dN/dS$ estimation largely infeasible: the controlled single-cell-type setting yields too few mutations per gene to generate statistically trustworthy selection coefficients, even for the subset of genes where values could technically be computed. We therefore present the following results purely as exploratory and do not interpret them directly, but note that the directional trends are broadly consistent with the *in vivo* findings.

With the above caveats in mind, selective pressure showed a directional correlation with ENU mutation burden ($r_{Spearman} = 0.362$, $p = 3.4\times10^{-22}$; Fig. 4a), consistent in direction with the *in vivo* pattern ($r = 0.262$), though the underlying gene-level $dN/dS$ estimates are insufficiently powered for direct interpretation. Genes with relatively low mutation rates tend to have smaller estimated $dN/dS$, whereas genes under the strongest positive selection accumulated higher mutation burdens. This enhanced signal in the controlled setting, where environmental and tissue heterogeneity are eliminated and many cell generations provide opportunity for both positive and negative selection, confirms that selection contributes independently to mutation patterns and is likely not an artifact of observational confounding, though further, more mutation dense experimentation is needed.

A GLM using expression, $dN/dS$, and degree explained 19% of variance in ENU-induced mutations, with selection contributing $\Delta R^2 = 0.168$, expression $\Delta R^2 = 0.021$, and degree contributing $\Delta R^2 = 0.006$ (Fig. 4c). This model was restricted to 421 genes with sufficient data for reliable parameter estimation. This reversal in relative contributions – selection now dominant, expression attenuated – is mechanistically revealing: it demonstrates that the two protective mechanisms operate at fundamentally different saturation thresholds. TCR is a molecular, rate-limited process that can be overwhelmed by sufficiently intense or repeated damage, while selection operates at the cellular population level across generations and is insensitive to damage rate. ENU therefore does not undermine the *in vivo* relevance of TCR – the persistent negative correlation between expression and mutation burden in ENU-treated cells confirms that transcription-coupled repair remains active – but rather dissociates the two mechanisms by saturating TCR while leaving selection intact. This controlled dissociation confirms that repair and selection are independent filters, and that their relative contributions to the observable mutation landscape depend on the intensity and chronicity of the mutagenic challenge.

As an independent validation of our experimental framework, we compared our ENU pathway-level results against those reported in [16], which applies a complementary analytical pipeline to the same ENU-treated fibroblast data using GO biological process terms. Briefly, [16] defines pathways using expressed genic and associated non-coding regulatory regions, using intronic sequences as the null to generate pathway scores, with the goal of estimating system-wide selection coefficients. We applied our framework to the same GO biological process gene sets to derive hyper- and hypo-mutated pathway scores. Despite differences in null model construction and scoring approach, pathway-level fold-change scores are significantly positively correlated across the two analyses (Pearson $r = 0.311$, $p = 5.55 \times 10^{-8}$; see Fig. 4d), confirming that the pattern of pathway-level protection and vulnerability we identify in the mutagenized data is not an artifact of our specific methodological choices but a robust signal recoverable by independent analyses.

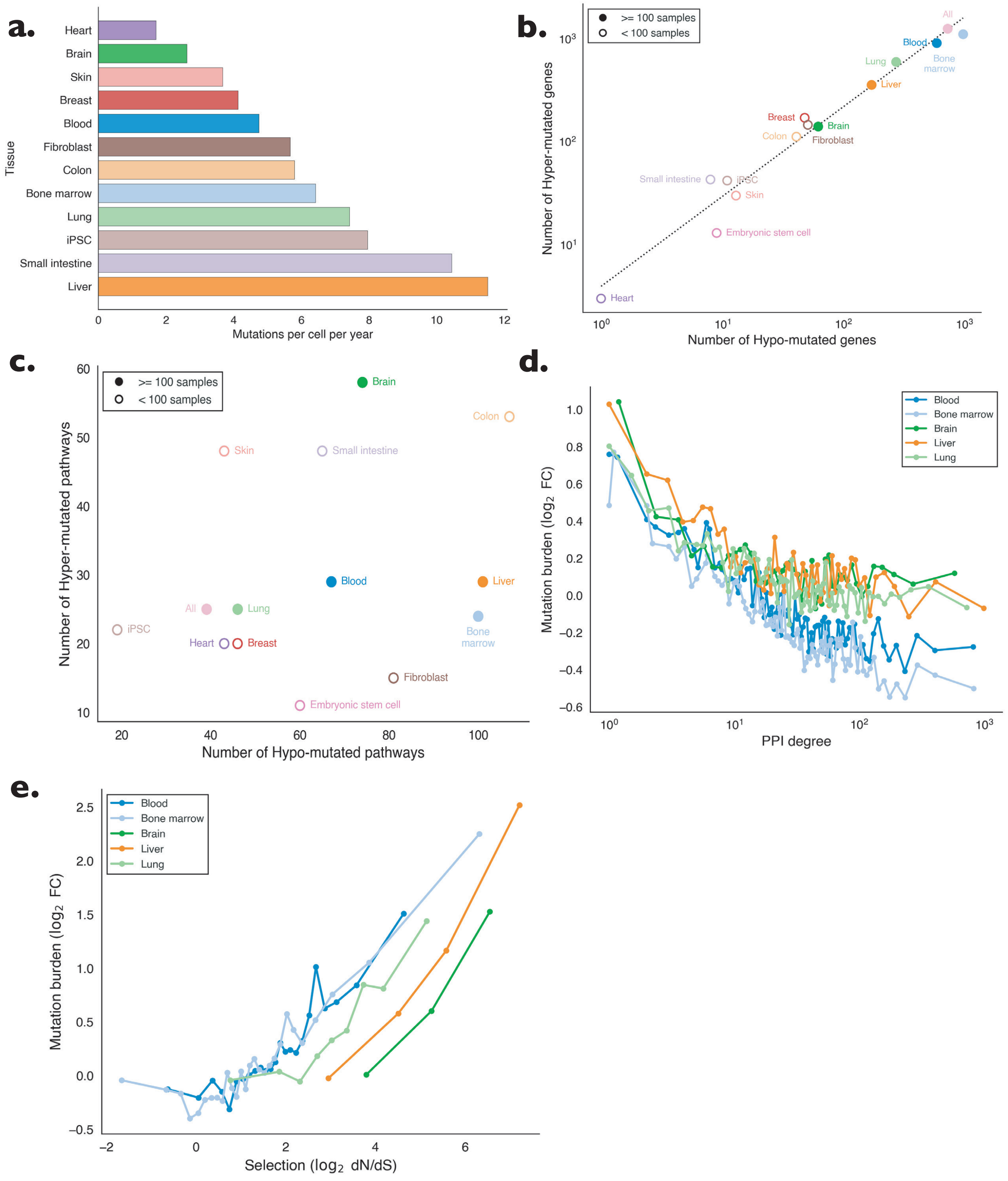


**Figure 1**: **Mutation burden across tissues.**

**Figure 1**: **Mutation burden across tissues. a)** Mutations per cell per year across different tissues. We observe an order of magnitude difference in the mutation rates of different tissues. **b)** Scatter plot showing the number of hypo-mutated genes versus the number of hyper-mutated genes for each tissue and for the aggregate dataset. Points are filled or open circles for tissues with $\geq$ 100 or $<$ 100 samples, respectively. **c)** The number of hypo-mutated pathways versus the number of hyper-mutated pathways across tissues. Points are filled or open circles for tissues with $\geq$ 100 or $<$ 100 samples, respectively. **d)** The relationship between human Interactome degree and gene-level mutation burden across tissues with $\geq$ 100 samples. All show a significant negative Spearman correlation, with blood and bone marrow exhibiting the strongest correlation (both $-0.20$), followed by liver and lung ($-0.09$ and $-0.10$), and, finally, brain ($-0.06$). **e)** The relationship between gene-level mutation burden and missense selection ($\log_2 dN/dS$) across tissues with $\geq$ 100 samples. All tissues are significantly positively correlated, ranging from a Spearman correlation of 0.25 (Bone marrow) to 0.44 (Liver).

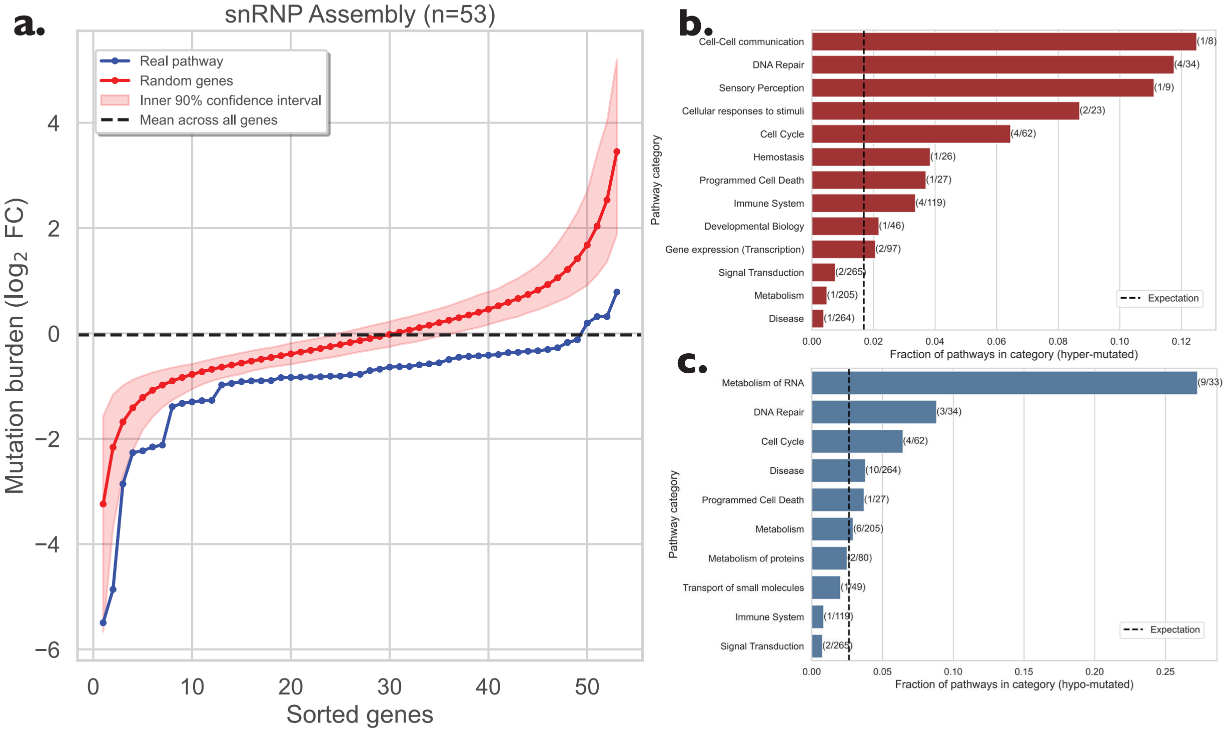


**Figure 2**: **Clustering of hyper- and hypo-mutated pathways into super categories. a)** Mutation profile of the snRNP assembly pathway, where the genes are sorted by mutation rate and plotted in blue next to a randomized version of the pathway in red with the same number of genes, where the shaded region is the 90th percentile of the null model. Even the most hyper-mutated genes in this do not exceed the average of the dataset, indicating that the hypo-mutated pathways avoid hyper-mutated genes. **b)** Hyper-mutated pathway super categories. Using pathway categorization based on Reactome, we count the number of hyper-mutated pathways in a category, and the total number of pathways in that category. Expected number of hyper-mutated pathways is represented by the dotted line, about 1.8% of pathways. Reactome categories with many hyper-mutated pathways appear at the top of the list. **c)** Hypo-mutated pathway super categories compared to expectation (2.6%). Exceptionally hypo-mutated categories appear at the top of the plot.

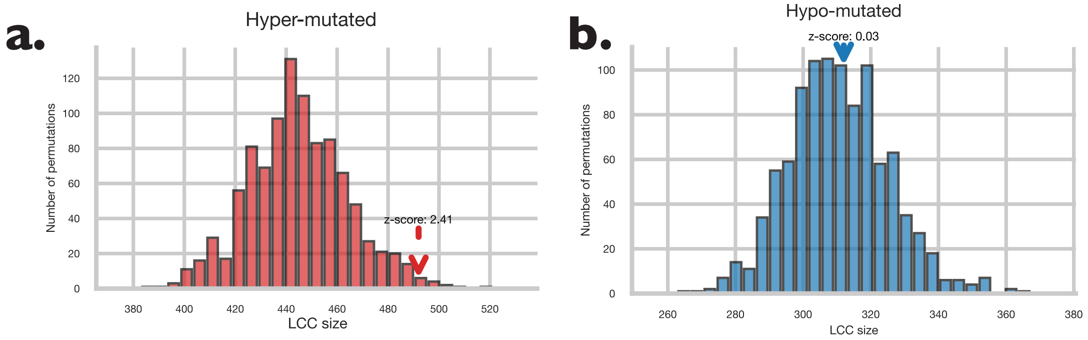


**Figure 3**: **Null distributions of the expected largest connected component sizes. a)** Hyper-mutated LCC under a degree-preserving randomization with 1,000 randomized gene sets; the arrow marks the observed LCC size (492; $z$-score = 2.41), indicating significant clustering of hyper-mutated genes. **b)** Hypo mutated LCC, with observed size (312; $z$-score $\approx$ 0.03). This is consistent with the null, reflecting that hypo-mutated genes are embedded in the existing core rather than forming a distinct peripheral module.

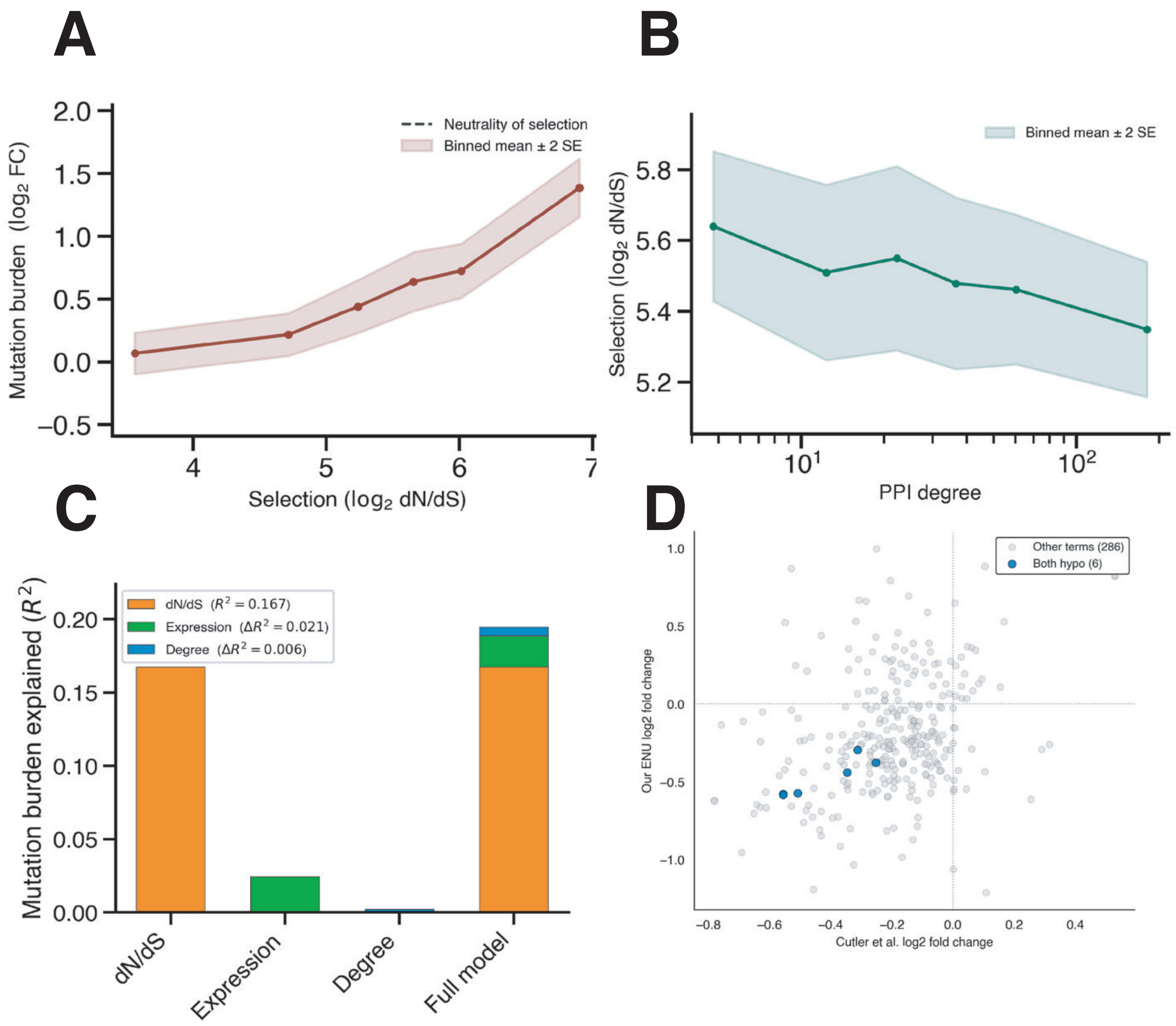


**Figure 4**: **ENU-mutagenized genetic mutations compared with aggregated *in vivo* mutations.** **a)** Selection ($\log_2(dN/dS)$) versus mutation burden in ENU-treated cells ($r_{Spearman} = 0.362$, $p = 3.4\times10^{-22}$), significantly stronger than the *in vivo* correlation ($r = 0.262$). **b)** Degree in the PPI versus $dN/dS$ in mutagenized samples, where we observe no significant correlation ($r_{Spearman} = -0.068$, $p = 0.0745$). **c)** Mutation burden varianced explained per variable in experimentally mutagenized samples. **d**) The $\log_2$ FC in our method vs. the method of [16] to estimate mutation burden of GO biological process gene sets.

### 7 Supplementary Data Captions

**Caption for Data S1. Gene-level mutation data observed versus expected mutation burden across tissues and ENU experiment.** Multi-sheet Excel table containing gene-level mutation burden results for the aggregated data, each individual tissue, and the ENU experimental dataset. Each sheet reports the gene identifier, expected number of mutations (via length-based null), observed mutation burden, $log_2$ fold change (observed/expected), and the nominal empirical p value.

**Caption for Data S2. Pathway-level mutation data observed versus expected mutation burden across tissues and ENU experiment.** Multi-sheet Excel table containing pathway-level mutation burden results in the same layout as the gene table, but with pathway identifiers (Reactome ID) in place of gene identifiers. Expected mutation burden and observed mutation burden have units of gene-level $\log_2$ FC.
